# Persistence of spore-forming *Clostridium* (*Clostridioides*) *difficile* through wastewater treatment plants in Western Australia

**DOI:** 10.1101/2022.10.19.512806

**Authors:** Jessica M. Chisholm, Papanin Putsathit, Thomas V. Riley, Su-Chen Lim

**Affiliations:** School of Biomedical Sciences, The University of Western Australia, Nedlands, WA, Australia; School of Medical and Health Sciences, Edith Cowan University, Joondalup, WA, Australia; Biosecurity and One Health Research Centre, Harry Butler Institute, Murdoch University, Murdoch, WA, Australia; PathWest Laboratory Medicine, Department of Microbiology, Nedlands, WA, Australia

**Keywords:** *Clostridium* (*Clostridioides*) *difficile*, molecular epidemiology, bacterial spore, wastewater, sewage, biosolids, One Health

## Abstract

There is growing evidence that shows *Clostridium* (*Clostridioides*) *difficile* is a pathogen of One Health importance with a complex dissemination pathway involving animals, humans and the environment. Thus, environmental discharge and agricultural recycling of human and animal waste have been suspected as factors behind the dissemination of *C. difficile* in the community. Here, the presence of *C. difficile* in 12 wastewater treatment plants (WWTPs) in Western Australia was investigated. Overall, *C. difficile* was found in 90.5% (114/126) of raw sewage influent, 48.1% (50/104) of treated effluent, 40% (2/5) of reclaimed irrigation water, 100% (38/38) of untreated biosolids, 95.2% (20/21) of anaerobically digested biosolids and 72.7% (8/11) of lime-amended biosolids. Over half the isolates (55.3%, 157/284) were toxigenic and 97 *C. difficile* ribotypes (RTs) were identified with RT014/020 the most common (14.8%, 42/284). Thirteen *C. difficile* isolates with the toxin profile A+B+CDT+ were found, including the hypervirulent RT078 strain. Resistance to the antimicrobials fidaxomicin, vancomycin, metronidazole, rifaximin, amoxicillin/clavulanate, meropenem and moxifloxacin was uncommon, however, resistance to clindamycin, erythromycin and tetracycline was relatively frequent at 56.7% (161/284), 14.4% (41/284) and 13.7% (39/284), respectively. This study revealed that toxigenic *C. difficile* was commonly encountered in WWTPs and being released into the environment. This raises concern about the possible spill-over of *C. difficile* into animal and/or human populations via land receiving the treated waste. In Western Australia, stringent measures are in place to mitigate the health and environmental risk of recycling human waste, however, further studies are needed to elucidate the public health significance of *C. difficile* surviving the treatment processes at WWTPs.

**IMPORTANCE:** *Clostridium difficile* infection (CDI) is a leading cause of antimicrobial-associated diarrhoea in healthcare facilities. Extended hospital stays and recurrences increase the cost of treatment, and morbidity and mortality. Community-associated CDI (CA-CDI) cases, with no history of antimicrobial use or exposure to healthcare settings, are increasing. The isolation of clinically important *C. difficile* strains from animals, rivers, soil, meat, vegetables, compost, treated wastewater and biosolids has been reported. The objective of this study was to characterise *C. difficile* in wastewater treatment plants (WWTPs) in Australia. We found that *C. difficile* can persist through the treatment processes of WWTPs and toxigenic *C. difficile* was being released into the environment becoming a potential source/reservoir for CA-CDI.

## INTRODUCTION

*Clostridium* (*Clostridioides*) *difficile* is an anaerobic Gram-positive spore-forming bacillus that causes disease ranging from uncomplicated diarrhoea to life-threatening pseudomembranous colitis, bowel perforation, sepsis and death (1). *C. difficile* is ranked as one of the most urgent antimicrobial resistance threats to public health by the United States Centers for Disease Control and Prevention (2, 3). Each year, *C. difficile* causes up to 500,000 infections and 29,000 deaths (∼6% mortality rate) in the United States (4). In Australia, *C. difficile* infection (CDI) results in approximately 8,500 cases and 500 deaths per year (5, 6), and is estimated to cost the healthcare system an average of AUD$12,704 per hospitalisation, a total of AUD $108 million per year (7).

The virulence of *C. difficile* is primarily attributed to toxins A and B, the genes for which are located on a pathogenicity locus (PaLoc) (8). An additional binary toxin (*C. difficile* transferase – CDT) encoded on the CDT locus (CdtLoc), but not found in all strains, was reported to be associated with severe disease (9). Traditionally, CDI has been considered a hospital-associated (HA) infection related to old age and antimicrobial exposure, especially to those antimicrobials with activity against the commensal gut flora which normally protects against overgrowth of *C. difficile* by inhibiting spore germination, vegetative growth and toxin production (1). Over the past two decades, the incidence of CDI has increased globally with rates remaining high in many high-income countries with growing reports of outbreaks (1, 10). Previously rare community-associated (CA) CDI, defined as cases with symptom onset in the community and no history of hospitalisation in the past 12 weeks or symptom onset ≤48 h of hospital admission (11), has emerged to represent a sizable proportion of CDI cases (1). In the United States, CA-CDI accounts for 41% of all CDI cases, an increase of four-fold from 1991 (12). In Australia, nearly 30% of all cases are CA, a four-fold increase from 1995 to 2011 (5, 13).

Using whole-genome sequencing (WGS), a study in the United Kingdom revealed that 45% of *C. difficile* strains isolated from 957 CDI cases were genetically distinct from all previous isolates (14). This refuted the traditional notion that CDI was primarily a HA infection and suggested that the transmission of *C. difficile* involved sources/reservoirs outside of the healthcare system. These are likely to be community-based, such as foodborne and/or environmental acquisition, as clinically important *C. difficile* strains have been isolated from animals, meat, vegetables, compost, gardens, lawns, rivers and lakes (15-17). This is in agreement with genomic studies in Europe and Australia that show (i) long-distance or cross-continental clonal transmission of *C. difficile* between humans and animals with ≤2 single nucleotide polymorphism (SNP) differences in their core-genome (18, 19), and (ii) genetically closely related *C. difficile* strains isolated from humans, food and the environment (retail potatoes, ready-to-eat salads, meat, compost, rivers, lakes, lawns and the end-products of wastewater treatment plants) (20-22). Taken together, this suggests a complex dissemination pathway of *C. difficile* between animals, humans and the environment. Thus, the environmental discharge and agricultural recycling of human and animal waste were hypothesized to be factors behind the widespread dissemination of *C. difficile* in the community which could lead to a rise in CDI.

Human sewage is treated through a series of physical, biological and chemical processes at wastewater treatment plants (WWTPs) to reduce organic matter and pathogen loads. However, the robustness of spore-forming pathogens such as Clostridia enables them to survive the treatment process with the potential to even flourish in the anaerobic digestive tanks commonly used to treat sludge (solid waste) from WWTPs. To date, few studies have investigated the presence and survival of *C. difficile* through WWTPs (23-26). The isolation of *C. difficile* strains in effluents (treated wastewater) has been described in Switzerland, Slovenia, England and New Zealand (23-26). In Canada, Xu *et al*. isolated *C. difficile* from effluents and biosolids as well as sediments taken from rivers connected to the effluent discharge pipes (27). The occurrence of *C. difficile* in WWTPs in Australia has not been explored. Thus, the objective of this study was to isolate and characterise *C. difficile* from influent (untreated wastewater), effluent, reclaimed irrigation water and biosolids from WWTPs in Western Australia (WA).

## MATERIALS AND METHODS

### Setting

In Australia, the processing of wastewater follows a specific sequence according to sewerage system guidelines (28). In WA, 80% of the wastewater collected is treated in the following ways: raw influent arriving at the WWTP goes through a preliminary treatment process which involves filtering the wastewater through screens and grit tanks to remove any large inorganic objects. The wastewater then flows into the primary sedimentation tank (also known as the settling tank or clarifier) where particles in the water gradually sink to the bottom of the tank to form sludge. The wastewater then flows into the aeration tank where microbes feed on oxygen, organic matter and any dissolved nutrients to produce carbon dioxide, nitrogen and more sludge. It then flows into the secondary sedimentation tank which is the final stage of treatment in most treatment plants before the treated effluent is released into the environment. In WA, the effluent is either returned to the ocean via large offshore pipes with small holes to ensure the effluent is evenly dispersed into the sea, used to replenish the groundwater or undergoes further advanced treatment before being recycled as irrigation water for sports grounds, public open spaces and non-food crops (e.g.: trees, turf and flowers), flushing the toilet, washing clothes, maintaining wetlands and industrial re-use of high-quality reclaimed water such as for use in the oil refining sector. The use of recycled water follows guidelines from the WA Department of Health (29). The sludge from WWTPs is either (i) collected and transferred to an anaerobic digester where organic matter is broken down into biogas and digested biosolids or (ii) dewatered using centrifugation with or without the addition of lime to reach a pH of >12. The management and subsequent use of biosolids follow guidelines from the WA Department of Water and Environment Regulation (previously known as the Department of Environment and Conservation) (30). These guidelines include a series of measures designed to manage the health and environmental risks associated with recycling biosolids, including determining the soil types, depth to groundwater, proximity to sensitive land and water resources, buffer distances to neighbouring protected areas, timing/seasons of application and slope of the land to prevent run-off (30).

### Sampling

A total of 126 influents, 104 effluents, 5 irrigation water samples and 70 biosolids (38 untreated, 21 anaerobically digested and 11 amended with lime) were collected from 12 WWTPs (W1 – W2) in WA, between January 2020 and June 2020 (**Table 1**). Influent and effluent samples were collected from all 12 WWTPs, biosolids from W5, W7 and W8 were untreated; anaerobically digested biosolids were from W2 and W12; lime-amended biosolids and reclaimed irrigation water were from W11.

**Table 1.**
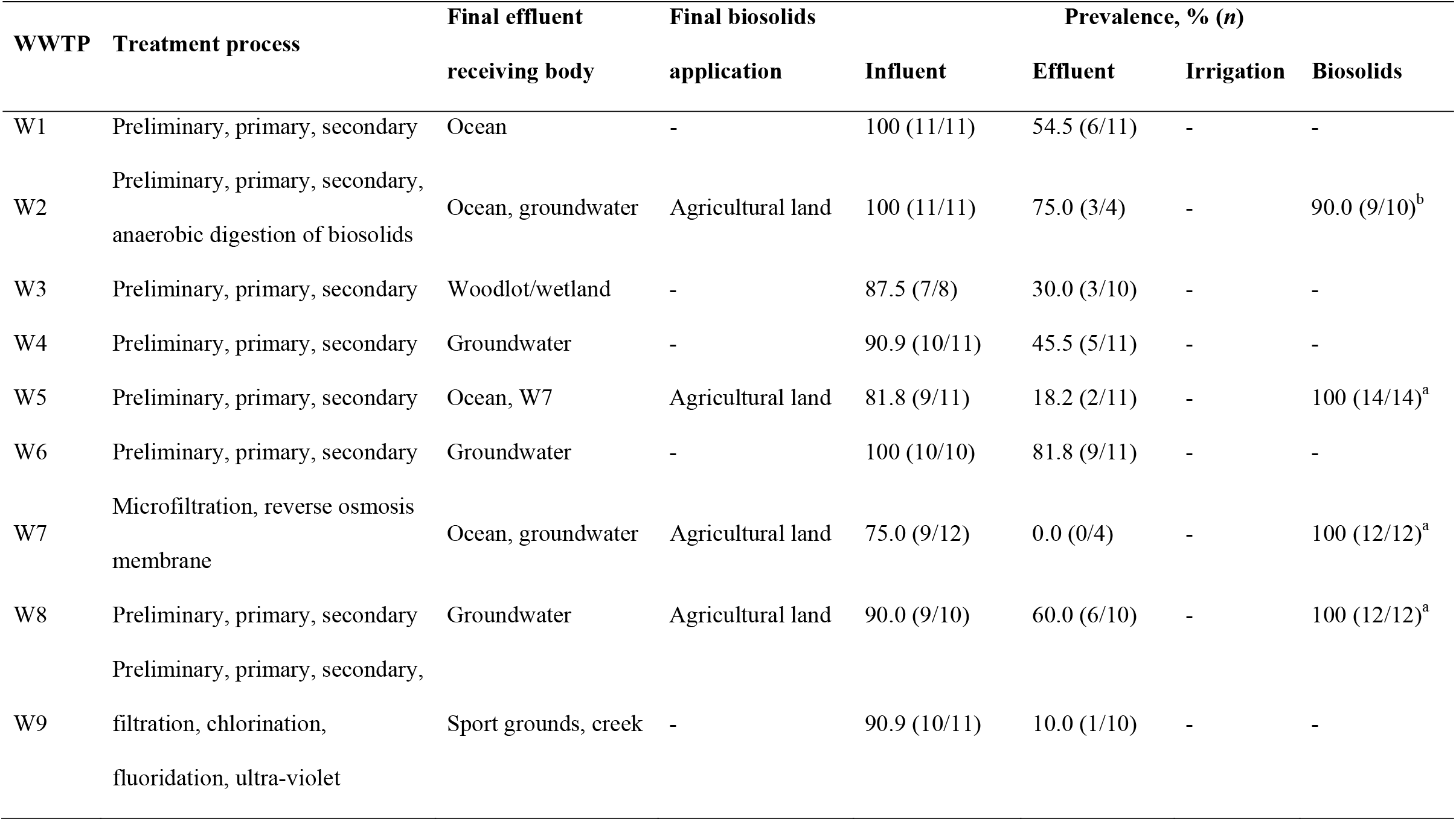
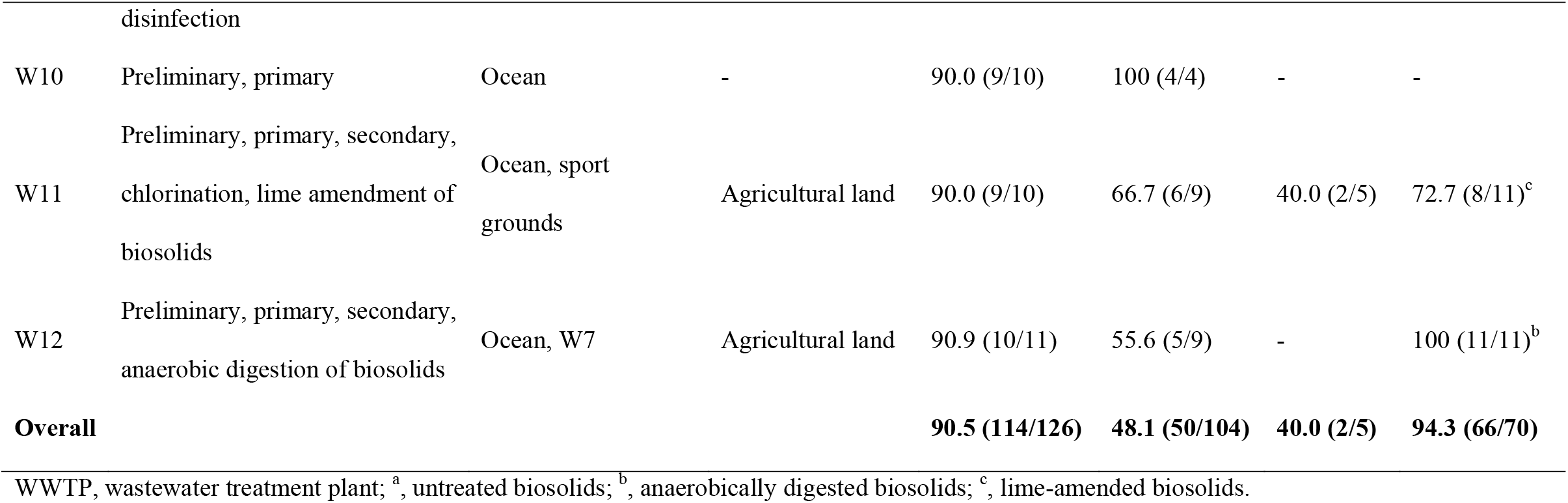
Characteristics of wastewater treatment plants and the prevalence of *C. difficile*.

| WWTP | Treatment process | Final effluent | Final biosolids | Prevalence, % (n) |  |  |  |
| --- | --- | --- | --- | --- | --- | --- | --- |
|  |  | receiving body | application | Influent | Effluent | Irrigation | Biosolids |
| W1 | Preliminary, primary, secondary | Ocean | - | 100 (11/11) | 54.5 (6/11) | - | - |
| W2 | Preliminary, primary, secondary, anaerobic digestion of biosolids | Ocean, groundwater | Agricultural land | 100 (11/11) | 75.0 (3/4) | - | 90.0 (9/10)<sup>b</sup> |
| W3 | Preliminary, primary, secondary | Woodlot/wetland | - | 87.5 (7/8) | 30.0 (3/10) | - | - |
| W4 | Preliminary, primary, secondary | Groundwater | - | 90.9 (10/11) | 45.5 (5/11) | - | - |
| W5 | Preliminary, primary, secondary | Ocean, W7 | Agricultural land | 81.8 (9/11) | 18.2 (2/11) | - | 100 (14/14)<sup>a</sup> |
| W6 | Preliminary, primary, secondary | Groundwater | - | 100 (10/10) | 81.8 (9/11) | - | - |
| W7 | Microfiltration, reverse osmosis membrane | Ocean, groundwater | Agricultural land | 75.0 (9/12) | 0.0 (0/4) | - | 100 (12/12)<sup>a</sup> |
| W8 | Preliminary, primary, secondary | Groundwater | Agricultural land | 90.0 (9/10) | 60.0 (6/10) | - | 100 (12/12)<sup>a</sup> |
| W9 | Preliminary, primary, secondary, filtration, chlorination, fluoridation, ultra-violet disinfection | Sport grounds, creek | - | 90.9 (10/11) | 10.0 (1/10) | - | - |
| W10 | Preliminary, primary | Ocean | - | 90.0 (9/10) | 100 (4/4) | - | - |
| W11 | Preliminary, primary, secondary,<br>chlorination, lime amendment of<br>biosolids | Ocean, sport<br>grounds | Agricultural land | 90.0 (9/10) | 66.7 (6/9) | 40.0 (2/5) | 72.7 (8/11)<sup>c</sup> |
| W12 | Preliminary, primary, secondary,<br>anaerobic digestion of biosolids | Ocean, W7 | Agricultural land | 90.9 (10/11) | 55.6 (5/9) | - | 100 (11/11)<sup>b</sup> |
| **Overall** |  |  |  | **90.5 (114/126)** | **48.1 (50/104)** | **40.0 (2/5)** | **94.3 (66/70)** |

WWTP, wastewater treatment plant; <sup>a</sup>, untreated biosolids; <sup>b</sup>, anaerobically digested biosolids; <sup>c</sup>, lime-amended biosolids.

### Culture conditions

Ten mL of each influent, effluent and reclaimed irrigation water sample was filtered through a 0.45 µm cellulose membrane (Pall Corporate, product ID 4761) using a manifold filtration system. The membrane was then placed in a 90 mL Brain Heart Infusion Broth supplemented with 5 g/L yeast extract, 1 g/L L-cysteine, 1 g/L taurocholic acid, 250 mg/L cycloserine and 8 mg/L cefoxitin (BHIB-S) (PathWest Media, Mt Claremont, WA) and incubated anaerobically in a Don Whitley Scientific Ltd (Otley, Yorkshire, United Kingdom) A35 anaerobic chamber (10% hydrogen, 10% carbon dioxide and 80% nitrogen) at 35°C for 7 days. Approximately 5 g of biosolid sample was added to 90 mL BHIB-S and incubated anaerobically as above. After incubation, 2 mL of broth was alcohol shocked by mixing an equal volume of absolute alcohol and standing for 1 h. The suspension was then centrifuged at 3,800 × *g* for 10 min and a 10 µl loop-full of pellet plated onto *C. difficile* ChromID agar (bioMérieux, France). The plates were incubated anaerobically and examined at 48 h. Unless there were colonies with different morphologies, one presumptive *C. difficile* colony per ChromID plate was sub-cultured onto a horse blood agar plate for identification based on colony morphology, odour and chartreuse fluorescence under UV light (∼360 nm) (31).

### Toxin profiling and ribotyping

Crude bacterial template DNA for toxin profiling was prepared by resuspension of cells in a 5% (wt/vol) solution of Chelex-100 resin (Sigma-Aldrich, Castle Hill, NSW, Australia). All isolates were screened by PCR for the presence of toxin A (*tcdA*), toxin B (*tcdB*) and binary toxin (*cdtA* and *cdtB*) genes (31). PCR ribotyping was performed as previously described (32). PCR ribotyping products were concentrated using a Qiagen MinElute PCR purification kit (Qiagen, Hilden, Germany). Visualisation of PCR products was performed with QIAxcel ScreenGel software (Qiagen, Hilden, Germany). Using the BioNumerics software package v.7.5 (Applied Maths, Saint-Martens-Latem, Belgium), the ribotyping patterns generated were compared to a reference library which consisted of over 16,000 *C. difficile* strains, including 54 internationally recognised RTs from the Anaerobe Reference Laboratory (ARL, Cardiff, United Kingdom) and the European Centre for Disease Prevention and Control (ECDC) collection. Isolates that gave patterns that did not correspond to any internationally recognised RTs in our library but had previously been isolated by our laboratory were assigned an internal nomenclature prefixed with QX. RTs that were new to our library were assigned a new QX number.

### Antimicrobial susceptibility testing

Minimum inhibitory concentrations (MICs) of a panel of 10 antimicrobial agents were determined by an agar incorporation method as described by the Clinical and Laboratory Standards Institute (CLSI) guidelines (33, 34). The panel included fidaxomicin, vancomycin, metronidazole, rifaximin, clindamycin, erythromycin, amoxicillin/clavulanic acid, moxifloxacin, meropenem and tetracycline. The clinical breakpoints for vancomycin and metronidazole were those recommended by the European Committee on Antimicrobial Susceptibility Testing (EUCAST) (35). The breakpoint for fidaxomicin of ≥1 mg/L was proposed by the European Medicines Agency (EMA) (36). Rifaximin resistance (≥32 mg/L) was as described (37), and the breakpoints for clindamycin, erythromycin, amoxicillin/clavulanic acid, moxifloxacin, meropenem and tetracycline were those provided by the CLSI (33).

### Statistical analysis

Fisher’s exact test was used, where appropriate, to compare the prevalence of *C. difficile* in samples from different WWTPs.

## RESULTS

### *C. difficile* prevalence

Overall *C. difficile* recovery was 90.5% (114/126) from influent, 48.1% (50/104) from effluent, 40% (2/5) from reclaimed irrigation water and 94.3% (66/70) from biosolids [100% (38/38) from untreated biosolids, 95.2% (20/21) from anaerobically digested biosolids and 72.7% (8/11) from lime-amended biosolids] (**Table 1**). Proportions of *C. difficile* recovered from influent collected at the 12 different WWTPs were not significantly different between sites (*p* = 0.78; range, 75% – 100%), but differences were detected in effluent (*p* = 0.004; range, 0% – 100%) and biosolids (*p* = 0.01; range, 75.7% – 100%) from different WWTPs.

### Toxin profiling and ribotyping of *C. difficile* isolates

Of the 284 *C. difficile* isolates recovered, 52.8% (150/284) contained *tcdA* and *tcdB* genes (A+B+), of which 8.7% (13/150) were also positive for binary toxin genes (*cdtA* and *cdtB*, CDT+). One hundred and twenty-seven isolates (44.7%) were non-toxigenic (A-B-CDT-). The remaining seven isolates yielded the following toxin profiles: A-B-CDT+ (1.4%, 4/284) and A-B+CDT+ (1.1%, 3/284). Toxigenic strains were isolated from 47.6% (60/126) of influent, 30.8% (32/104) of effluent, 71.4% (50/70) of biosolids and none of the five irrigation water samples.

Ninety-seven *C. difficile* RTs were identified, 37 (38.1%) of which were internationally recognised, 43 (44.3%) were classified with internal nomenclature and 17 (17.5%) were novel (**Figure 1**). The majority of the novel strains (64.7%, 11/17) were non-toxigenic. The most common RT found was RT014/020 (A+B+CDT-) which comprised 14.8% (42/284) of the isolates, followed by RT010 (A-B-CDT-) (14.4%, 41/284). The 13 *C. difficile* strains with toxin profile A+B+CDT+ were RT078 (*n* = 7), RT126 (3), RT127 (2) and QX 656 (1). Toxigenic *C. difficile* RT014/020 represented the majority of isolates found in influent (12.5%, 16/128), effluent (25.5%, 13/51) and biosolids (12.6%, 13/103) but not in irrigation water (0%, 0/2).

**Figure 1.**
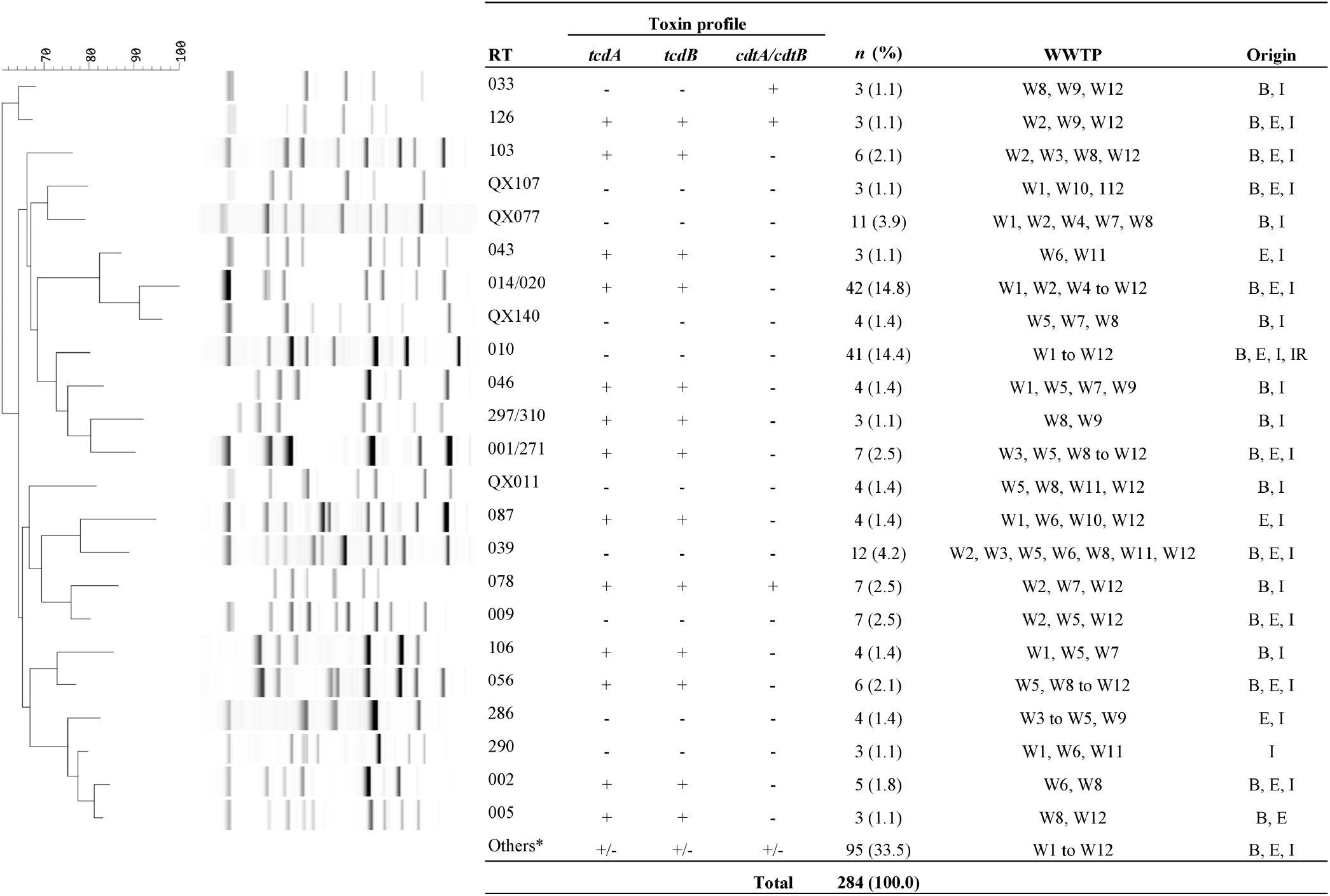
Summary of *C. difficile* ribotyping patterns and toxin gene profiles of isolates obtained from wastewater treatment plant in WA. Ribotyping pattern analysis was performed with a neighbour-joining tree using Pearson correlation. WWTP, wastewater treatment plant; B, biosolids; E, effluents; I, influents; IR, reclaimed irrigation water. *RTs with a frequency of two or fewer. These were RTs 003, 012, 015/193, 018, 026/118, 051, 053, 064, 070, 081, 101, 125, 127, 137, 247, 281, 584, 605 and local RTs (17 singletons, QXs 002, 024, 025, 026, 029, 042, 051, 057, 068, 076, 086, 087, 099, 121, 141, 161, 189, 195, 210, 211, 229, 243, 365, 372, 395, 488, 491, 509, 532, 541, 605, 610, 629, 656, 657, 658, 659 and 660).

### Antimicrobial susceptibility of *C. difficile* isolates

Most isolates were susceptible to the antimicrobials recommended for treating CDI (**Table 2**). First-line treatment drugs, vancomycin and metronidazole inhibited 98.2% and 100% of the isolates, respectively. The in vitro activity of rifaximin was the most potent (MIC50/MIC90, 0.03/0.03 mg/L), followed by fidaxomicin (MIC50/MIC90, 0.25/0.25 mg/L), metronidazole (MIC50/MIC90, 0.25/0.5 mg/L) and vancomycin (MIC50/MIC90, 2/2 mg/L). All isolates were susceptible to fidaxomicin, metronidazole and amoxicillin/clavulanic acid. The proportions of isolates resistant to vancomycin (1.8%, 5/284), rifaximin (1.4%, 4/284), meropenem (0.35%, 1/284) and moxifloxacin (2.5%, 7/284) were low. Forty-one isolates (14.4%) were resistant to erythromycin, all with an MIC of >256 mg/L. These isolates were RT039 (*n* = 6), RT014/020 (4), QX011 (4), RT046 (3), RT009 (2), RT078 (2), RT126 (2), RT247 (2), QX024 (2), QX395 (2), RT001/271 (1), RT005 (1), RT010 (1), RT051 (1), RT053 (1), RT056 (1), RT127 (1), QX099 (1), QX140 (1), QX658 (1) and two novel RTs. Resistance to tetracycline was observed in 13.7% (39/284) of the isolates, including RT078 (*n* = 6), RT014/020 (4), RT039 (4), RT046 (3), QX011 (3), RT010 (2), RT126 (2), RT247 (2), QX395 (2), RT005 (1), RT009 (1), RT012 (1), RT106 (1), RT127 (1), QX024 (1), QX107 (1), QX141 (1), QX658 (1) and two novel strains. Over half the isolates (56.7%, 161/284) were resistant to clindamycin. Among the frequently isolated RTs, resistance to clindamycin was most common in the non-toxigenic RT039 (91.7%, 11/12) with a geometric mean (GM) of 27.4 mg/L compared to 6.75 mg/L of all isolates. *C. difficile* RT039, a non-toxigenic strain also, displayed greater resistance to tetracycline (GM of 3.36 mg/L vs 0.56 mg/L of all isolates).

**Table 2.** Antimicrobial susceptibility data for *C. difficile* isolates, by RT.

| RT | Antimicrobial | MIC range | MIC50/MIC90 | GM | S |  | I |  | R |  |
| --- | --- | --- | --- | --- | --- | --- | --- | --- | --- | --- |
|  |  |  |  |  | % | *n* | % | *n* | % | *n* |
| All RTs<br>(*n* = 284) | FDX | 0.03-0.5 | 0.25/0.25 | 0.16 | 100.0 | 284 | - | - | - | - |
|  | VAN | 1-4 | 2/2 | 1.45 | 98.2 | 279 | - | - | 1.8 | 5 |
|  | MTZ | 0.125-1 | 0.25/0.5 | 0.26 | 100.0 | 284 | - | - | - | - |
|  | RFX | 0.015 to >64 | 0.03/0.03 | 0.03 | 98.6 | 280 | - | - | 1.4 | 4 |
|  | CLI | 0.125 to >256 | 8/32 | 6.75 | 12.3 | 35 | 31.0 | 88 | 56.7 | 161 |
|  | ERY | 0.25 to >256 | 2/>256 | 1.51 | 85.6 | 243 | - | - | 14.4 | 41 |
|  | AMC | 0.125-2 | 0.5/1 | 0.61 | 100.0 | 284 | - | - | - | - |
|  | MEM | 1-16 | 2/4 | 2.34 | 98.6 | 280 | 1.1 | 3 | 0.35 | 1 |
|  | MXF | 1-32 | 2/2 | 2.06 | 94.7 | 269 | 2.8 | 8 | 2.5 | 7 |
|  | TET | 0.125-64 | 0.25/16 | 0.56 | 83.8 | 238 | 2.5 | 7 | 13.7 | 39 |
| RT 014/020<br>(*n* = 42) | FDX | 0.03-0.5 | 0.25/0.25 | 1.7 | 100.0 | 42 | - | - | - | - |
|  | VAN | 1-2 | 1/2 | 1.37 | 100.0 | 42 | - | - | - | - |
|  | MTZ | 0.125-0.5 | 0.25/0.25 | 0.25 | 100.0 | 42 | - | - | - | - |
|  | RFX | 0.015-16 | 0.03/0.03 | 0.03 | 100.0 | 42 | - | - | - | - |
|  | CLI | 0.5 to >256 | 8/16 | 6.61 | 9.5 | 4 | 23.8 | 10 | 66.7 | 28 |
|  | ERY | 0.25 to >256 | 2/4 | 1.58 | 90.5 | 38 | - | - | 9.5 | 4 |
|  | AMC | 0.25-1 | 0.5/0.5 | 0.51 | 100.0 | 42 | - | - | - | - |
|  | MEM | 1-4 | 2/2 | 2.07 | 100.0 | 42 | - | - | - | - |
|  | MXF | 2-2 | 2/2 | 2.00 | 100.0 | 42 | - | - | - | - |
|  | TET | 0.125-64 | 0.25/1 | 0.42 | 90.5 | 38 | - | - | 9.5 | 4 |
| RT 010<br>(*n* = 41) | FDX | 0.03-0.5 | 0.25/0.5 | 0.21 | 100.0 | 41 | - | - | - | - |
|  | VAN | 1-2 | 1/2 | 1.31 | 100.0 | 41 | - | - | - | - |
|  | MTZ | 0.25-0.5 | 0.25/0.25 | 0.26 | 100.0 | 41 | - | - | - | - |
|  | RFX | 0.015-0.03 | 0.03/0.03 | 0.03 | 100.0 | 41 | - | - | - | - |
|  | CLI | 1-64 | 8/16 | 6.42 | 9.8 | 4 | 36.6 | 15 | 53.7 | 22 |
|  | ERY | 1 to >256 | 2/2 | 1.74 | 97.6 | 40 | - | - | 2.4 | 1 |
|  | AMC | 0.5-1 | 1/1 | 0.95 | 100.0 | 41 | - | - | - | - |
|  | MEM | 2-4 | 2/4 | 2.76 | 100.0 | 41 | - | - | - | - |
|  | MXF | 2-4 | 2/2 | 2.07 | 95.1 | 39 | 4.9 | 2 | - | - |
|  | TET | 0.125-32 | 0.25/0.5 | 0.31 | 95.1 | 39 | - | - | 4.9 | 2 |
| RT 039<br>(*n* = 12) | FDX | 0.125-0.5 | 0.25/0.25 | 0.22 | 100.0 | 12 | - | - | - | - |
|  | VAN | 1-2 | 1/2 | 1.33 | 100.0 | 12 | - | - | - | - |
|  | MTZ | 0.25-0.5 | 0.5/0.5 | 0.42 | 100.0 | 12 | - | - | - | - |
|  | RFX | 0.015-0.03 | 0.03/0.03 | 0.03 | 100.0 | 12 | - | - | - | - |
|  | CLI | 4 to >256 | 16/>256 | 27.43 | - | - | 8.3 | 1 | 91.7 | 11 |
|  | ERY | 2 to >256 | 2/>256 | 2.00 | 50.0 | 6 | - | - | 50.0 | 6 |
|  | AMC | 0.5-1 | 1/1 | 0.94 | 100.0 | 12 | - | - | - | - |
|  | MEM | 2-4 | 2/4 | 2.67 | 100.0 | 12 | - | - | - | - |
|  | MXF | 2-2 | 2/2 | 2.00 | 100.0 | 12 | - | - | - | - |
|  | TET | 0.25-16 | 2/16 | 3.36 | 66.7 | 8 | - | - | 33.3 | 4 |
| Eighty-eight<br>'other' RTs<br>(*n* = 189) | FDX | 0.03-0.5 | 0.25/0.25 | 0.15 | 100.0 | 189 | - | - | - | - |
|  | VAN | 1-4 | 2/2 | 1.51 | 97.4 | 184 | - | - | 2.6 | 5 |
|  | MTZ | 0.125-1 | 0.25/0.5 | 0.25 | 100.0 | 189 | - | - | - | - |
|  | RFX | 0.015 to >64 | 0.03/0.03 | 0.03 | 97.9 | 185 | - | - | 2.1 | 4 |
|  | CLI | 0.125 to >256 | 8/32 | 6.40 | 14.3 | 27 | 32.8 | 62 | 52.9 | 100 |
|  | ERY | 0.25 to >256 | 2/>256 | 1.43 | 84.1 | 159 | - | - | 15.9 | 30 |
|  | AMC | 0.125-2 | 0.5/1 | 0.57 | 100.0 | 189 | - | - | - | - |
|  | MEM | 1-16 | 2/4 | 2.31 | 97.9 | 185 | 1.6 | 3 | 0.5 | 1 |
|  | MXF | 1-32 | 2/2 | 2.07 | 93.1 | 176 | 3.2 | 6 | 3.7 | 7 |
|  | TET | 0.125-64 | 0.25/16 | 0.61 | 81.0 | 153 | 3.7 | 7 | 15.3 | 29 |

FDX, fidaxomicin; VAN, vancomycin; MTZ, metronidazole; RFX, rifaximin; CLI, clindamycin; ERY, erythromycin; AMC,
amoxicillin/clavulanic acid; MEM, meropenem; MXF, moxifloxacin; TET, tetracycline; MIC, minimum inhibition concentration; GM,
geometric mean; S, susceptible; I, intermediate; R, resistant. MICs are given in mg/L.

Multidrug-resistance (MDR), defined as resistance to ≥3 antimicrobial classes, was observed among 7.0% (20/284) of *C. difficile* isolates. These were RT046 (*n* = 3), QX011 (3), RT014/020 (2), RT078 (2), RT005 (1), RT039 (1), RT051 (1), RT126 (1), RT127 (1), RT247 (1), QX024 (1), QX395 (1), QX658 (1) and one novel strain. The most common MDR profiles were resistance to clindamycin, erythromycin and tetracycline (85.0%, 17/20). Two isolates, RT005 and QX011, were resistant to five antimicrobials (rifaximin, clindamycin, erythromycin, moxifloxacin and tetracycline). Two isolates (one RT126 and one QX011) were resistant to clindamycin, erythromycin, moxifloxacin and tetracycline; and a single RT046 strain was resistant to rifaximin, clindamycin, erythromycin and tetracycline.

## DISCUSSION

In this study, we found that toxigenic *C. difficile* was commonly encountered in WWTPs and was being released into the environment, consistent with earlier European studies. In Switzerland, Romano *et al*. isolated *C. difficile* in 100% of their influent and treated effluent samples from nine WWTPs with 85% of the strains being toxigenic (26). This raised concerns about the possible contamination of local rivers that were receiving the treated effluents and the safety of reusing treated effluents. In 2015, Steyer *et al*. performed a 1-year survey on the occurrence of enteric pathogens in effluent from a conventional two-stage activated sludge WWTP in Slovenia (23). Astroviruses were not found in August and September, and hepatitis A and E viruses were not detected at all, while rotaviruses, noroviruses, sapoviruses and *C. difficile* were detected in all samples collected throughout the study period. In total, 121 *C. difficile* strains with 32 different RTs were isolated in the study, of which RT014/020 and RT010 were the most prevalent. This confirmed the authors’ previous findings of *C. difficile* being widely distributed in Slovenian rivers, including RT014, with a positive correlation with increased population densities (38). Thus, the release of effluent into local rivers was suspected of being a potential source of *C. difficile* contamination. In 2018, 186 *C. difficile* strains isolated from effluent samples from 18 WWTPs in England were sequenced and compared with 70 clinical isolates using phylogenetic analysis (24). *C. difficile* from human clinical cases and WWTPs were genetically highly related. Overall, these studies confirmed the extensive release of toxigenic *C. difficile* into surface waters and question the need to improve strategies for the removal of bacterial spores associated with causing human diseases from the effluent before release into the environment.

In comparison, our study yielded a similarly high prevalence of *C. difficile* in influent (90.5%) but the overall isolation of *C. difficile* in effluent was lower at 48.1%. This was likely due to the inclusion of WWTPs that performed tertiary (e.g., sand filtration, chlorination) and advanced water (e.g., membrane filtration, reverse osmosis) treatment. All effluent samples from W7, a WWTP that uses membrane filtration and reverse osmosis to produce high quality reclaimed water, did not contain *C. difficile*, and only 10% of effluent from W9 had *C. difficile* likely attributed to the plant performing membrane filtration, chlorination and ultra-violet disinfection in addition to the traditional activated sludge treatment process. Effluents from the remaining WWTPs had a *C. difficile* prevalence ranging between 18.2% to 100%, averaging 58.7%. However, this finding should be interpreted with some care given the small number of samples collected from each WWTP. While the release of effluent into water bodies can lead to contamination and therefore potentially contribute to CA-CDI in at-risk populations upon exposure to environmental *C. difficile* spores, we suspect the risk of acquiring CDI via effluent discharged into the ocean is low in WA as, in our recent study, *C. difficile* was not found in any of the 89 seawater samples collected along the coast of WA near Perth, the State capital (39). Furthermore, the use of effluent as irrigation water is not widespread in WA with only a small number of local councils using it to irrigate their sport grounds and public spaces. However, a notable proportion of our effluent was used to replenish the groundwater which is commonly extracted to irrigate golf courses, public spaces and residential lawns. The presence and bacterial load of *C. difficile* in groundwater, as well as fields that use groundwater as irrigation, was not studied here and hence requires further investigation.

In Canada, Xu *et al*. reported a high prevalence of *C. difficile* in raw sludge (92%), anaerobically digested sludge (96%), lime-amended biosolids (73%) and sediments (39%) from rivers that received the effluent that has been chlorinated, passed through sand filters and de-chlorinated prior to disposal into rivers (27). This is comparable with our findings of *C. difficile* in 100% of untreated biosolids, 95.2% of anaerobically digested biosolids and 72.7% of lime-amended biosolids. In Australia, the management of biosolids is strictly regulated, with a pathogen (P1-P4) and chemical contaminant (C1-C3) grading determining their suitability for different end uses (30). The highest quality biosolids (P1C1) have no restrictions in application, and biosolids with a P1C2 grading can be used in land application for crops that may be consumed raw. Mid-quality biosolids (P2C2 and P3C2) can be used for landscaping, horticulture, forestry and pasture, and crops that are processed before consumption and have no direct contact with soil. Low-quality biosolids with a P4C3 grading are destined for composting or landfill. The application of biosolids with such a high prevalence of *C. difficile* may lead to widespread dissemination of *C. difficile* in the environment. However, none of the biosolids tested in this study was of the highest quality (P1C1) with unrestricted use as the biosolids produced by the WWTPs in WA were predominantly mid-quality with a P2C2 or P3C2 grade. For biosolids to be graded as P2 or P3, they must undergo treatment such as (i) composting according to the Australian Standard AS 4454 which is reaching an internal temperature of 55°C for 3 consecutive days with a minimum of three turns for lower risk materials such as plant and vegetation, and 15 days with a minimum of five turns for high-risk materials, or (ii) the addition of lime so that the pH is maintained at >12 for 72 h, and the effectiveness of these treatments at reducing pathogen levels must be validated by testing the level of faecal indicator organisms (*Escherichia coli* and viable helminths) (30). In this study, although the level of *C. difficile* was not known, the survival of *C. difficile* in biosolids treated with lime was seen with a 72.7% prevalence. In addition, composting is insufficient at reducing *C. difficile* spores to an undetectable level (40, 41). This was attributed to *C. difficile* being a spore-former capable of withstanding high temperatures and harsh environmental conditions (41). In fact, *C. difficile* has been isolated in commercially composted products in Australia and the United States at a prevalence of 22.5% and 35.9%, respectively (42, 43). In WA, a conservative approach was adopted to protect the environment from nutrient run-off and reduce the risk of exposure to the general public. Currently, biosolids are no longer applied to forestry or pasture, and are only supplied to agriculture land where crops will be processed before consumption and have no direct contact with the soil. Thus, risk of exposure to *C. difficile* via biosolids is presumably low. Nevertheless, the dissemination of *C. difficile* via commercial compost is likely, and worth investigating. The length of time that *C. difficile* spores can persist on pasture and in biosolids-incorporated soil also remains to be determined.

In agreement with Moradigaravand *et al*., we found an overlap between *C. difficile* genotypes isolated from WWTPs and those isolated from humans in the same area, with 65 out of 97 RTs found in WWTPs also isolated from hospital patients (24). Having been isolated from influent, effluent and biosolids across 11 WWTPs, *C. difficile* RT014 was the most common among all strains (14.8%). *C. difficile* RT014 is a lineage of One Health importance, well-established in humans, pigs and the environment in Australia (19). Nationwide, *C. difficile* RT014 accounts for approximately 30% of all CDI cases in humans and 23% of isolates from neonatal pigs (44, 45). In WA, *C. difficile* RT014 also represents a significant proportion of *C. difficile* isolates from lawn (39%), compost (10%), hospital gardens (14%), home gardens (21%), root vegetables (7%) and water bodies (11%) (31, 39, 42, 46-48). Although the zoonotic and environmental transmission of *C. difficile* is still being debated, recent data suggest long-distance transmission of *C. difficile* between animals, environmental reservoirs and humans (18, 19, 39, 42, 49, 50). In our recent study where 142 *C. difficile* RT014 strains from humans, animals, compost, hospital gardens, lawns, root vegetables, shoes and water bodies were sequenced and analysed, extensive co-clustering of human, animal and environmental strains was found, with 9% of the human strains having a clonal relationship (≤2 SNP) indicative of direct transmission and 60% closely related (≤9 SNP) to at least one animal or environmental strains (22). We suspect the land application of biosolids and animal manure has aided in the dissemination of *C. difficile* in the environment and may have contributed to a rise in CA-CDI. In addition, it is worth noting that *C. difficile* RT078 isolated from the biosolids of W2, W7 and W12, is a hypervirulent strain associated with severe CDI in a younger population and more frequently CA in Europe (51). It was the third most common RT of *C. difficile* isolated from CDI patients in 34 European countries (52). Even though human CDI cases caused by RT078 are uncommon in Australia (44), the finding of RT078 in biosolids is a particular concern from a public health standpoint.

Not surprisingly, the antimicrobial resistance (AMR) profile of *C. difficile* from WWTPs in Australia was similar to those from humans; with no or low level resistance to fidaxomicin, vancomycin, metronidazole, rifaximin, amoxicillin/clavulanate, meropenem and moxifloxacin, but high level resistance to clindamycin (53, 54). Interestingly, the non-toxigenic *C. difficile* RT039 displayed a high level of resistance to clindamycin and tetracycline. Due to RT039 being non-toxigenic, it is usually not reported from human clinical samples (53, 54), however, it is a relatively common RT found in the environment in Australia although greater resistance to any antimicrobial agents has never been reported (55). Non-toxigenic *C. difficile* strains are highly prevalent in Asia and many of these strains are resistant to multiple antimicrobials, possibly due to inappropriate antimicrobial use in the region (56). These *C. difficile* RT039 strains may be of Asian origin and could have either acquired AMR genes via horizontal gene transfer from other microbes in the host intestine or from the consortium of pathogenic/commensal microbes in WWTPs. The presence of diverse selective pressures in a WWTP could create favourable conditions for the transfer of AMR genes and the proliferation of AMR bacteria. AMR is a growing global health threat, yet environmental surveillance of AMR is largely lacking and WWTPs could potentially be a focal point in the fight against AMR.

Although a low number of samples per sample type (influent, effluent, biosolids and reclaimed water) was collected from each WWTPs and quantification of *C. difficile* was not performed, the importance of effluent and biosolids in *C. difficile* release into the environment was clearly demonstrated in our study. In summary, we showed that WWTP influent, effluent and biosolids in WA contained toxigenic *C. difficile* strains belonging to RTs found in human CDI cases. Thus, the release of effluent and biosolids can create a source or reservoir of community infection. Future studies are needed to determine the public health significance of *C. difficile* surviving the treatment processes at WWTPs and commercial composting processes. With climate change and increased water scarcity, the development of efficient wastewater treatment and recycling practices is imperative to securing a sustainable future. WWTPs should have a central role in environmental surveillance of AMR and emerging infectious diseases, and this will require a multi-disciplinary One Health approach.

## ACKNOWLEDGEMENTS

We are grateful to the staff of the Water Corporation (WA) for providing advice, and for collecting samples.

